# A deep learning framework for high-throughput mechanism-driven phenotype compound screening

**DOI:** 10.1101/2020.07.19.211235

**Authors:** Thai-Hoang Pham, Yue Qiu, Jucheng Zeng, Lei Xie, Ping Zhang

## Abstract

Target-based high-throughput compound screening dominates conventional one-drug-one-gene drug discovery process. However, the readout from the chemical modulation of a single protein is poorly correlated with phenotypic response of organism, leading to high failure rate in drug development. Chemical-induced gene expression profile provides an attractive solution to phenotype-based screening. However, the use of such data is currently limited by their sparseness, unreliability, and relatively low throughput. Several methods have been proposed to impute missing values for gene expression datasets. However, few existing methods can perform *de novo* chemical compound screening. In this study, we propose a mechanism-driven neural network-based method named DeepCE (Deep Chemical Expression) which utilizes graph convolutional neural network to learn chemical representation and multi-head attention mechanism to model chemical substructure-gene and gene-gene feature associations. In addition, we propose a novel data augmentation method which extracts useful information from unreliable experiments in L1000 dataset. The experimental results show that DeepCE achieves the superior performances not only in *de novo* chemical setting but also in traditional imputation setting compared to state-of-the-art baselines for the prediction of chemical-induced gene expression. We further verify the effectiveness of gene expression profiles generated from DeepCE by comparing them with gene expression profiles in L1000 dataset for downstream classification tasks including drug-target and disease predictions. To demonstrate the value of DeepCE, we apply it to patient-specific drug repurposing of COVID-19 for the first time, and generate novel lead compounds consistent with clinical evidences. Thus, DeepCE provides a potentially powerful framework for robust predictive modeling by utilizing noisy omics data as well as screening novel chemicals for the modulation of systemic response to disease.

---

Target-based high-throughput screening dominates conventional drug discovery process which follows a one-drug-one-gene paradigm. It has been the focus of computer-aided drug discovery for decades including recent application of deep learning. However, the readout from the modulation of a single protein by a chemical is poorly correlated with organism-level therapeutic effect or side effect. As a result, the failure rate from a lead compound generated from the target-based screening to approved drug is high. Phenotype-based screening has created renewed interests for identifying cell-active compounds but suffered from low-throughput and difficulty in target deconvolution. Therefore, a high-throughput, mechanism-driven phenotype compound screening method will no doubt facilitate drug discovery and development.

Gene expression profiling has been widely used to characterize cellular and organismal phenotypes. Systematic analysis of genome-wide gene expression of chemical perturbations on human cell lines has led to significant improvements in drug discovery and systems pharmacology. In particular, it can be applied to drug repurposing^1,1–4^, discovering drug mechanisms^5^, lead identification^6^, and predicting side effects for pre-clinical compounds^7^. The use of genome-wide chemical-induced gene expression was initially made possible by the appearance of Connectivity Map (CMap)^8^, which consists of gene expression profiles of five human cancer cell lines perturbed by ~ 1300 compounds after 6h. However, the limited data availability across cell types restricts the performances of these analyses which heavily depend on the coverage of chemicals and human cell lines. To overcome this limitation, a novel gene expression profiling method, L1000, which is the extension of CMap project, was developed by NIH library of integrated network-based cellular signatures (LINCS) program^9^. After Phase I of LINCS program, L1000 dataset consists of ~ 1,400,000 gene expression profiles on the responses of ~ 50 human cell lines to one of ~ 20,000 compounds across a range of concentrations. Recently, L1000 dataset and its normalization versions^10^ are widely used in drug repurposing and discovery^11,12^. Despite these successes, there are several major problems when utilizing L1000 dataset.

First, although the number of gene expression profiles is much larger than that in CMap, many missing expression values remain in the vast combinatorial space of chemicals and cell lines. Second, there are hundreds of millions of drug-like purchasable chemicals which are potential drug candidates^13^. It is infeasible to experimentally test all of these chemicals for their chemical-induced gene expression profiles across multiple cell lines. Finally, due to various experimental problems (e.g. batch effect), many experiment measurements are not reliable (as shown in Supplementary Figure 1). These serious obstacles will limit the effectiveness and scope of utilizing L1000 dataset in drug discovery. Therefore, predicting gene expression values for unmeasured and unreliable experiments are necessary.

Missing entries in the combinatorial space is the problem of not only L1000 dataset but also other gene expression profiling datasets. Before the appearance of L1000 dataset, several methods of imputing missing values have been proposed for gene expression datasets. We categorize these methods into two main approaches depending on the dependence of other information besides gene expression data. The first approach does not use any additional information. Works following this approach include k nearest neighbor (kNN)^14^, singular value decomposition^14^, least mean square^15–17^, Bayesian principal component analysis^18^, Gaussian mixture clustering^19^, and support vector regression^20^. The second approach uses additional information to predict expression profiles. For example, chemical structures are used to predict chemical-induced gene expressions but that work does not consider cell-specific information^21^.

The approaches described above are designed for matrix-structured data (i.e. gene × experiment) while L1000 dataset is formulated as tensor-structured data (i.e. gene × chemical × cell × doses × time) so they cannot be applied to capture high-dimensional associations that help to impute missing values for L1000 dataset. Recently, several methods are proposed to predict gene expression profiles in L1000 dataset. In particular, to deal with high-dimensional structured data, an extension of linear regression model named polyadic regression is developed to capture interactions emerging across features^22^. Matrix completion methods are also adapted to handle tensor-structured gene expression data^23,24^.

Above methods for L1000 dataset just focus on imputing the missing values of some gene expression profiles or the whole gene expression profiles of some missing experiments. They are not very useful in the real setting of drug discovery where the chemical-induced gene expression profile of new chemicals needs to be identified. This motivates us to solve a more practical but more challenging problem: predicting gene expression profiles for *de novo* chemicals (i.e. chemicals that do not appear in training data). Solving this problem is necessary because it helps to infer gene expression profiles of new chemicals without conducting experiments that require time and human resources. More importantly, this problem can be expanded to predict gene expression profiles for new cell lines which can be difficult for measuring in *in vitro* environment. However, current computational approaches for predicting gene expression values for L1000 dataset cannot work well in *de novo* setting. In particular, tensor completion approach cannot predict gene expression profiles for new chemicals because of the inaccessibility to chemical features. Polyadic regression, theoretically, can predict gene expression profiles for high-dimensional data in *de novo* chemical setting because of using chemical features. However, in practice, it is not feasible because of huge computational resources required for handling high-dimensional data (i.e. this method fails when applied to more than 3-dimensional data). Therefore, there is a strong incentive to develop a new and effective method that exploits high-dimensional data for predicting gene expression profiles for de novo chemical setting.

To address the aforementioned problems, we design a mechanism-driven neural network-based model, DeepCE, which captures high-dimensional associations among biological features as well as non-linear relationships between biological features and outputs to predict gene expression profiles given a new chemical compound. Our proposed DeepCE significantly outperforms the state-of-the-art models for predicting gene expression profiles in L1000 dataset not only for *de novo* chemical setting but also for traditional imputation setting. Several novelties in the architecture of model contribute to the success of DeepCE. First, we leverage graph convolutional network to automatically extract chemical substructure features from data. Second, attention mechanism is used to capture associations among chemical substructures and genes, and among genes in cell lines. Finally, gene expression values of all L1000 genes are predicted simultaneously from hidden features by multi-output multi-layer feed-forward neural network. Besides developing this neural network-based model, we propose a data augmentation method by which we can extract useful information from unreliable experiments in L1000 dataset to improve the prediction performance of our model. We also verify the effectiveness of DeepCE by comparing the performances of several classification models trained on gene expression profiles generated from DeepCE and those trained on original gene expression profiles in L1000 dataset for two downstream tasks: drug-target and disease predictions. Finally, we assess the value of our proposed method for the challenge and urgent problem, finding treatment for COVID-19, by *in silico* screening all chemical compounds in Drugbank against COVID-19 patient clinical phenotypes. The prioritized lead compounds are consistent with existing clinical evidences. To our knowledge, it is the first work of phenotype-based drug repurposing for COVID-19. The source code of DeepCE and the generated gene expression profiles of all chemical compounds in Drugbank are publicly available for research purpose, which could make significant a contribution to drug discovery and development in particular, and computational chemistry and biology research in general.

## Methods

In this section, we present datasets used in our study and our proposed model, DeepCE, as well as baseline models for predicting gene expression profiles including linear models, vanilla neural network, k-nearest neighbor, and tensor-train weight optimization models. The general framework of training and testing these computational models for L1000 gene expression profile prediction is shown in Figure 1. Basically, computational models take L1000 experimental information (i.e. chemical compound, cell line, time stamp, and chemical dose size) from L1000 dataset as inputs, transform them into numerical representations, and then predict L1000 gene expression profiles based on these representations. The details of the numerical feature transformation process for chemical and biological objects used in our study and model implementation of DeepCE and other baselines are shown in Supplementary Notes. Moreover, in this section, we present the data augmentation method that extracts useful information from unreliable experiment in L1000 dataset to improve the prediction performance of our models, and the evaluation method for our models.

**Figure 1.**
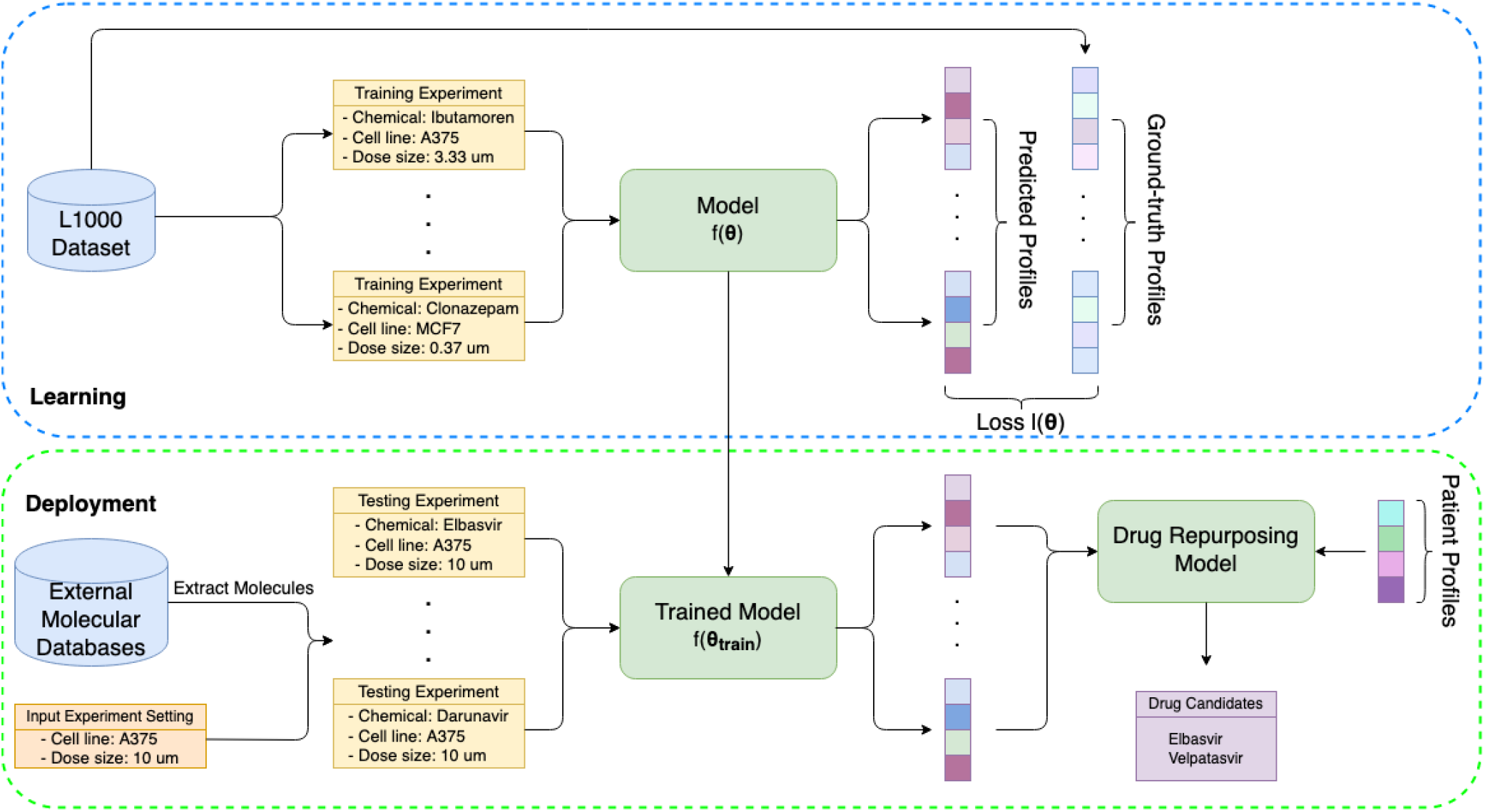
General framework of training computational models for L1000 gene expression profile prediction and using them for downstream application (i.e. drug repurposing). The objective for the learning process is minimizing the loss between predicted profiles and grouth-truth profiles in L1000 dataset. After training, models is used for generating profiles for new chemicals in external molecular database (e.g. DrugBank, ChEMBL). These profiles are then used for *in silico* screening to find potential drugs for disease treatment

### Datasets

#### Bayesian-based peak deconvolution L1000 dataset

After the original version of L1000 dataset was released9, many efforts have been made to improve the quality of this dataset. For example, instead of using k-means clustering algorithm as in the original version, some works propose to use Gaussian mixture model to enhance the accuracy of peak deconvolution step^25,26^. One work, in another way, develops a multivariate method called Characteristic Direction to compute gene signatures instead of using the moderated Z-score as in the original version^10^. In our study, we conduct experiments on Bayesian-based peak deconvolution L1000 dataset which has been shown to generate more robust z-score profiles from L1000 assay data, and therefore, gives better representation for perturbagens^27^. In particular, we train and evaluate our proposed methods on level 5 data of this dataset. The gene expression profiles result from experiments of 7 most frequent cell lines and 6 most frequent chemical dose sizes in L1000 dataset are used to construct our gene expression dataset. We then select high-quality experiments from our dataset and split into high-quality training set, and development and testing set. We also construct the original training set by keeping unreliable experiments in our gene expression dataset and the augmented training set generated by our data augmented algorithm. The details of constructing these sets are described in Supplementary Notes. The statistics of these training, development, and testing sets are shown in Supplementary Table 1.

#### STRING database for human protein-protein interactions

STRING^28^ is a multi-source database of protein-protein interactions. These interactions which can be known or predicted, direct (physical) or indirect (functional) are collected from five main sources including genomic context prediction, high-throughput lab experiments, conserved co-expression, automated text-mining, and previous knowledge databases. In our setting, we extract the human protein-protein interaction network (i.e. ~ 19,000 nodes (proteins) and ~ 12,000,000 edges (interactions)) from this database to compute vector representations for L1000 genes. The drug-target vector representations for chemical compounds used in our study are also computed from this human protein-protein interaction network. The details of generating these representations from STRING database are shown in Supplementary Notes.

#### Drugbank database for drug-target interaction and disease predictions

Drugbank is a well-known, comprehensive database used in many bioinformatics and cheminformatics tasks^29^. This database consists of information about drugs and their targets. In our experiments, we extract ATC labels derived from the first level of ATC tree and targets of drugs appeared in L1000 dataset from Drugbank. There are 698 drug targets and 14 ATC labels in the extracted dataset. We select the most frequent ATC labels and drug targets based on their frequents on this dataset as the labels of drugs to form drug-target prediction and ATC prediction datasets. These datasets are used to evaluate the performance of gene expression profiles generated from our models.

#### Patient expression in response to SARS-CoV-2 infection

Patient expression data for this study is downloaded from NCBI Gene Expression Omnibus(GSE147507).^30^ We used expression profile from SARS-CoV-2 patient and healthy negative controls in series 15 for differential expression analysis. Two technical replicates are from one male diseased patient (age 74) and 2 uninfected samples are from one male (age 72) and one female (age 60). DESeq2^31^ package is used to generate the differential gene expression profile of the patient. Not all L1000 genes appear in the result of DESeq2 package so we only consider genes appear in both L1000 dataset and DESeq2 package (i.e. 838 genes) when comparing with chemical-induced gene expression profiles.

### Overall architecture of DeepCE

Our neural network-based model for L1000 gene expression profile prediction, DeepCE, consists of several components as follows. First, we use graph convolutional network to learn numerical representation for chemical compound from its graph structure and feed-forward neural network to learn numerical representations for cell line and chemical dose size. We also use numerical representations for L1000 genes which are derived from the human protein-protein interaction network (described in Supplementary Notes). After that, these vector representations are put into the interaction component to capture high-level feature associations including chemical substructure-gene and gene-gene feature associations. Finally, the prediction component takes the outputs of the interaction component as inputs to predict the gene expression values for all L1000 genes simultaneously. The overall architecture of DeepCE and its hyperparameters used in our experiments are shown in Figure 2 and Supplementary Table 3 respectively. The following paragraphs describe each component of DeepCE in detail.

**Figure 2.**
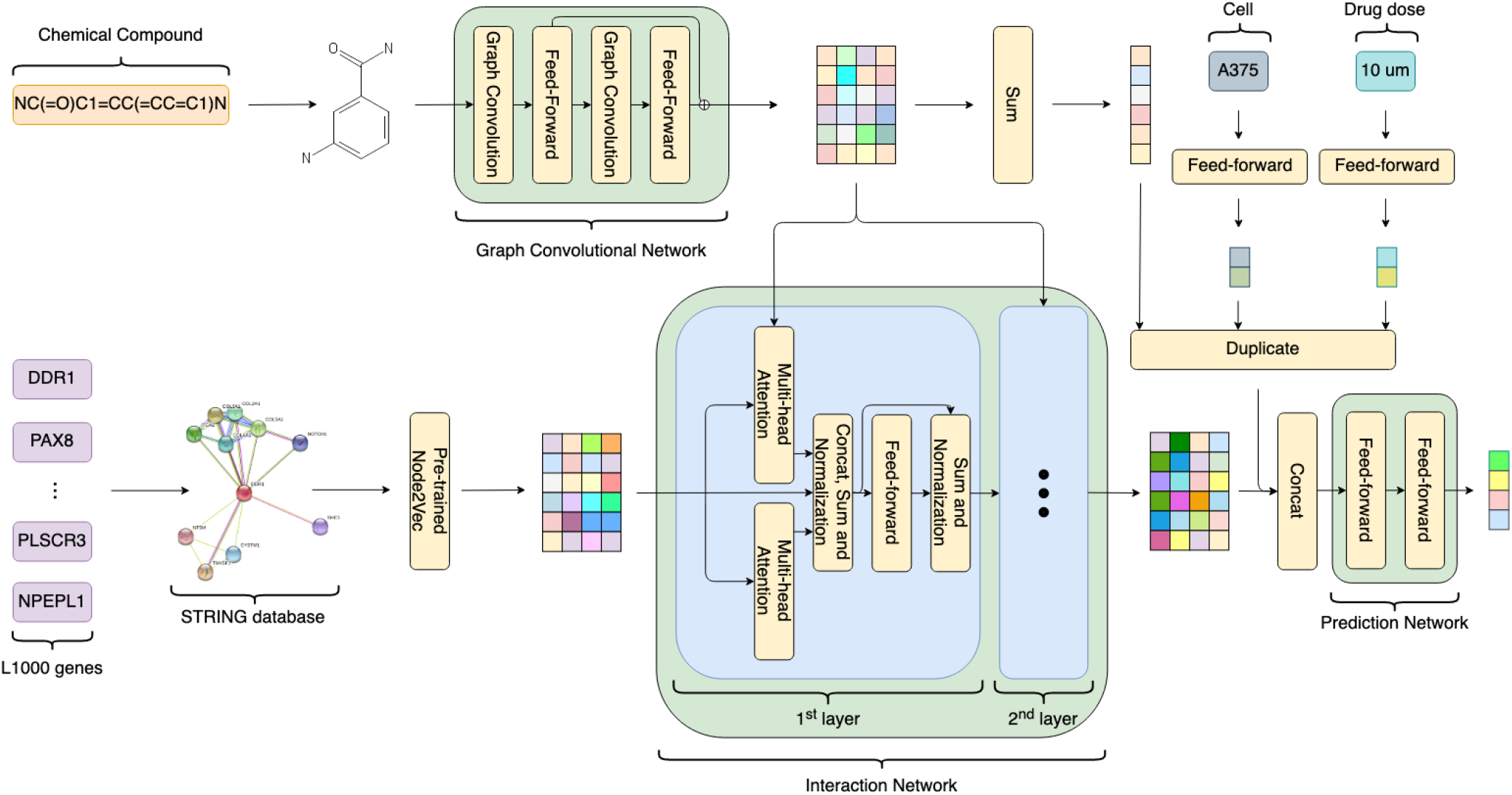
Overall architecture of DeepCE (The details of 2^*nd*^ layer which has similar architecture to the 1^*st*^ layer in the interaction network are omitted to save space)

#### Graph convolutional network for neural fingerprint

Recently, data-driven chemical fingerprints are shown to be more effective than predefined chemical fingerprints (e.g. PubChem, ECFP) for many biological prediction problems. Therefore, we propose to use graph convolutional network (GCN) to capture the chemical substructure information. The original GCN model for chemical fingerprint32 takes a graph structure of chemical compound as input and update vector representations for each node (atom) in graph (chemical compound) from its neighborhoods by convolutional operation. Thus, the vector for each node after convolutional operation can be seen as the representation of chemical substructures. The final vector which is the sum of vectors of every node is used as the chemical fingerprint. GCN model used in our experiments is primarily based on that model but with a minor modification. In particular, we output vector representations for every nodes instead of one vector representation for the chemical compound because we want to model the associations of chemical substructure features with gene features. In our settings, we use the GCN model with 2 convolutional layers (radius R = 2). It means that the output vector from GCN for each atom represents the chemical substructure which is a span of 2-hop distance from that atom. The initial representations for atoms and bonds are multi-hot vectors that capture the symbol, degree, number of Hydro neighborhoods, and aromaticity of atoms, and type of bonds that have lengths of 62 and 6, respectively. The details of GCN model used in our experiments are shown in Supplementary Algorithm 1.

#### Multi-head attention for gene-gene and chemical substructure-gene feature associations

Attention mechanisms where an element of one set selectively focuses on a subset of another set (attention) or its set (self-attention) based on attention weights are used widely in neural network-based models and effectively applied to many AI tasks including computer vision and natural language processing. In our experiments, we propose to apply the attention method named multi-head attention for modeling associations among gene features, and among gene and chemical substructure features. Multi-head attention was first proposed in Transformer model which achieves state-of-the-art results for many natural language processing tasks^33^. Basically, each element in sets can be represented by a set of three vectors query, key, and value. An individual attention module is a function of mapping queries and sets of key-value pairs to output matrix computed by:

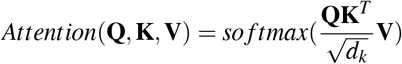

where **Q, K, V** are matrices (sets) of queries, keys, values respectively and *d_k_* is a scaling factor. Multi-head attention focuses on different representation subspaces by concatenating several individual attention modules:

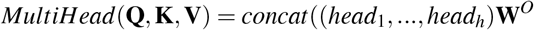

where 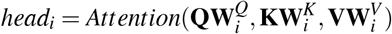.

This multi-head attention mechanism is the main ingredient used to construct the interaction component of DeepCE. In particular, the interaction component consists of two identical layers where outputs of the first layer are used as inputs for the second layer. For each layer, we use 2 separate multi-head attention modules with 4 heads for each module to model associations among genes in gene set and among elements in gene set and chemical substructure set. Length of query, key, value vectors is set at 512. Outputs from these two multi-head attention modules are concatenated and put into normalization layer followed by feed-forward layer and another normalization layer. The abstract architecture of interaction component is shown in Figure 2.

#### Multi-output prediction

The multi-output prediction component which is a 2-layer feed-forward neural network with ReLU activation function takes input as the concatenation of chemical neural fingerprint, gene feature generated by interaction component, cell line and chemical dose size features to predict gene expression values for all L1000 genes together as follows:

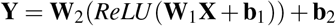

where **W**_1_, **W**_2_, **b**_1_, **b**_2_ are weight matrices and bias vectors of this network. The output size of this feed-forward neural network is set at 978 which is the number of L1000 genes.

#### Objective function

The objective function used in DeepCE model is mean squared error (MSE) between predicted and ground-truth gene expression values and is computed as follows:

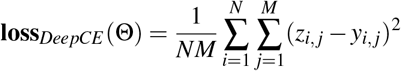

where **Θ** are the set of parameters in DeepCE model. *N* and *M* the number of gene expression profiles in a batch and number of L1000 genes respectively. *z_i, j_* and *y_i,j_* are ground-truth and predicted gene expression values of *j^th^* gene in *i^th^* gene expression profile.

### Baseline Models

In this section, we describe several baseline models used in our experiments including linear models, vanilla neural network, k-nearest neighbor, and tensor-train weight optimization^24^.

#### Linear models

We experiment with multi-output linear regression model and its regularization versions including Lasso regression (L1 regularization) and ridge regression (L2 regularization) models. Like DeepCE, input for these models is the concatenation of numerical representations for chemical, gene, cell line, and chemical dose size features but we use predefined chemical fingerprints and drug-target features instead of data-driven representations derived from GCN for chemicals. The details of these representations are described in Supplementary Notes. Multi-output linear models can be seen as 1-layer feed-forward neural network without activation function.

#### Vanilla neural network

The vanilla neural network used in our experiments can be seen as the simpler version of DeepCE model that does not include the interaction network component for modeling gene-gene and gene-chemical substructure feature associations and GCN for generating neural fingerprints. Input for this vanilla neural network is similar to its for linear models. The following layers in this network are similar to the prediction network component in DeepCE model which is a 2-layer feed-forward neural network with ReLU activation function.

#### K-nearest neighbor

We also propose a k-nearest neighbor-based approach for gene expression prediction for *de novo* chemical setting. In particular, gene expression profile for a new chemical compound in one particular setting (i.e. cell line, chemical dose size) is generated by averaging gene expression profiles of its nearest neighborhoods in the training set in the same setting. In our research, we experiment with different numbers of neighborhoods from 1 to 15 and different distance measures including cosine, euclidean, correlation, Jaccard and Tanimoto distances.

#### Tensor-train weight optimization

Tensor-train weight optimization (TT-WOPT) is a tensor completion approach proposed to retrieve missing values in tensor data from existing values. It has been shown to be effective for predicting missing values of L1000 dataset which can be formulated as a tensor-structure object without using additional information^24^. In our research, we conduct experiments to compare it with our proposed model, especially in *de novo* chemical setting. Because this model does not require additional information so input for it is L1000 gene expression values formulated as a tensor.

### Data augmentation

From Supplementary Figure 1, we can see that only a small number of experiments in L1000 dataset are reliable (i.e. APC score ≥ 0.7) so it would be wasteful if we cannot exploit useful information from a large number of unreliable experiments. It will be shown in the Results section (i.e. Table 1) that simply adding unreliable experiments to the high-quality training set (original training set) makes the performances of our models worse. Thus, we propose the data augmentation method by which we can effectively exploit unreliable experiments to improve the performances of our models. We argue that although an experiment (level 5 data) is unreliable, not all its bio-replicates experiments (level 4 data) are also unreliable and we will extract these reliable bio-replicate experiments by our proposed data augmentation method. The basic idea is that we, first, train our model on the high-quality training set, and then, generate predicted gene expression profiles for unreliable experiments. These predicted gene expression profiles are compared with their bio-replicate gene expression profiles and we incorporate bio-replicate gene expression profiles that have the similarity scores with their predicted gene expression profiles larger than the threshold. Supplementary Algorithm 2 presents this data augmentation method in detail. In our settings, the similarity score is Pearson correlation.

**Table 1.**
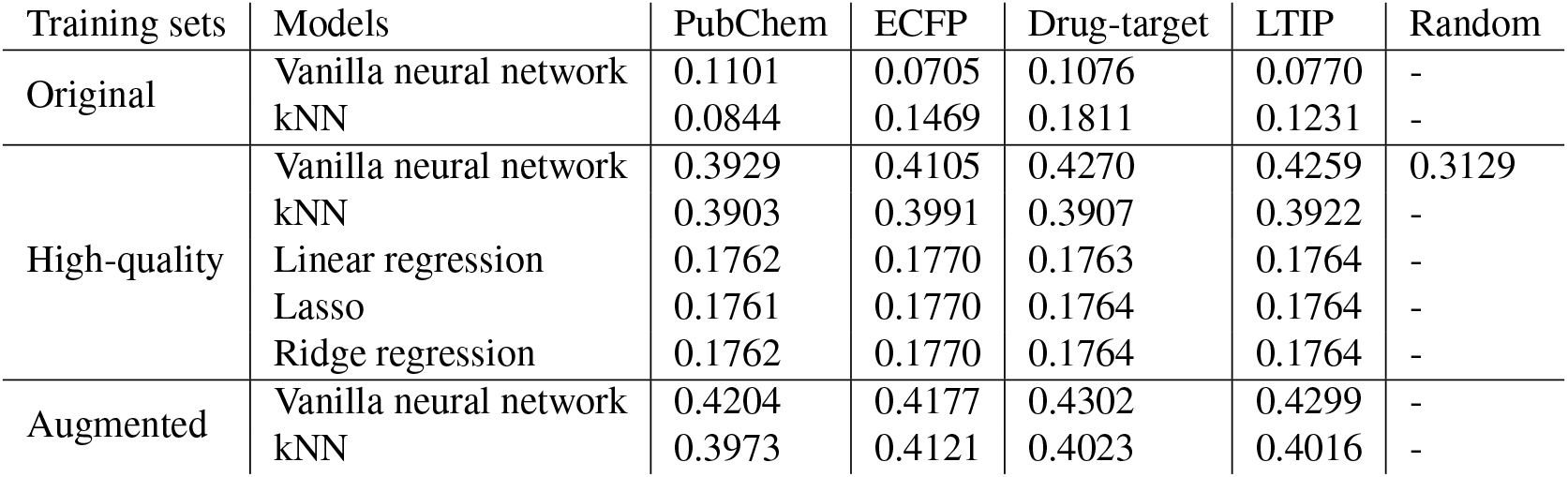
Performances (Pearson correlation) on testing set of vanilla neural network, kNN, and linear models with different chemical features trained with different training sets

### Performance evaluation

Pearson correlation coefficient is used to evaluate performances of models in our experiments. Correlation scores which measure the relationship between ground-truth and predicted gene expression profiles have been shown to be more effective than error measures for microarray data analysis^34,35^. Moreover, using Pearson correlation allows us to conduct unbiased evaluation for our models which are optimized for mean squared error. We calculate the average Pearson correlation for a dataset as follows:

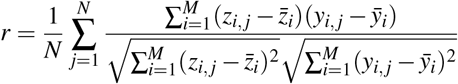

where N and M are number of gene expression profiles in the dataset and number of L1000 genes respectively. 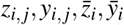 are ground-truth and predicted gene expression values of *j^th^* gene in *i^th^* gene expression profile and ground-truth and predicted mean values of *i^th^* gene expression profile.

Besides using Pearson correlation to directly evaluate the quality of our predicted gene expression profiles, we also use area under the receiver operating characteristic curve (AUC) to verify the effectiveness of these predicted profiles for downstream binary classification tasks including drug-target and ATC code predictions.

## Results and Discussions

### DeepCE significantly outperforms baseline models in the novel chemical setting

In this experiment, we compare DeepCE model and its simpler variants constructed by removing either the whole interaction component or just one part of its (i.e. chemical substructure-gene or gene-gene feature association modules) with several baseline models including vanilla neural network, kNN, linear models, and TT-WOPT. While TT-WOPT predicts output based on gene expression values only, other models learn the relationship between experimental information and gene expression profiles to make predictions. For DeepCE, we use neural fingerprints while for other models, we use predefined fingerprints including PubChem and circular (ECFP6) fingerprints, and drug-target information including latent target interaction profile (LTIP)^36^ and our proposed drug-target feature to represent chemicals. All models are trained on the high-quality training set and are evaluated on the test set.

As listed in Table 1 and Table 2, DeepCE model and its variants achieve order-of-magnitude improvements over baseline models. In particular, DeepCE model significantly outperforms other models including vanilla neural network, kNN, linear models, and TT-WOPT by achieving a Pearson correlation of 0.4907 on the testing set (paired t-test, *p – value* < 4.63 × 10^−15^). Comparing to its simpler variants whose interaction components are removed, DeepCE also achieves better performance, indicating that the effectiveness of modeling chemical substructure-gene and genegene feature associations. Specifically, the performance of DeepCE decreases to 0.4620, 0.4477, and 0.4418 when removing chemical substructure-gene feature association part, gene-gene feature association part, and the whole interaction component (paired t-test, *p – value* < 2.25 × 10^−5^), respectively. For baseline models, vanilla neural networks and kNN achieve pretty good performances. Linear models including linear regression, Lasso, and ridge regression do not work well for our problem. It indicates that the linear relationship is not sufficient to model the dependencies among variables in this dataset. TT-WOPT, which does not leverage additional features besides gene expression values to make the predictions, as we expect, does not work well for *de novo* chemical setting. In particular, it achieves a Pearson correlation of 0.0144 which is similar to randomness.

**Table 2.** Performances (Pearson correlation) on testing set of TT-WOPT and DeepCE with its simpler variants trained with different training sets

| Training sets | Models | Performances |
| --- | --- | --- |
| High-quality | TT-WOPT | 0.0133 |
|  | DeepCE w/o interaction component | 0.4418 |
|  | DeepCE w/o chemical substructure-gene attention | 0.4620 |
|  | DeepCE w/o gene-gene attention | 0.4477 |
|  | DeepCE | 0.4907 |
| Augmented | DeepCE | 0.5014 |

### DeepCE outperforms the state-of-the-art methods in the imputation setting

We further investigate the performance of DeepCE for the traditional imputation setting that does not require chemicals in the testing set to be different from chemicals in the training set, and compare it with TT-WOPT which has been shown to be effective for this setting. To do that, we randomly split the high-quality dataset to the new training, development, and testing sets and conduct the experiment on these sets. Note that, at this time, we split the dataset by gene expression profile instead of chemical compound. The details of the training, development, and testing set for imputation setting are shown in Supplementary Table 2.

For the traditional imputation setting, we observe DeepCE outperforms TT-WOPT with a large margin. In particular, DeepCE achieves a Pearson correlation of 0.7010 compared to its of 0.5113 of TT-WOPT. This result indicates that DeepCE consistently achieves the best performances for both *de novo* chemical and traditional imputation settings by effectively leveraging features of chemical and biological objects including chemical compounds and genes.

### Chemical similarity has an impact on prediction performance

To investigate thoroughly the prediction performance of our models, we investigate the impact of chemical similarity between testing set and training set. In particular, we compute the distance between one experiment in the testing set and its nearest neighbors experiments in the training set which are induced by the most similar chemicals (i.e. determined by comparing their fingerprints with the fingerprint of the chemical compound induced the experiment in the testing set) on the same cell line. The distance between the two experiments is the Tanimoto coefficient of PubChem fingerprints of their two chemicals, and the distance between the experiment on the testing with its nearest neighbor experiments in the training set is the average of distances between that experiment and each of its nearest neighbors. After computing the distances to the training set for all experiments on the testing set, we sort them by the ascending order and compare the Pearson correlation scores of these experiments. We calculate the average Pearson correlation scores of all experiments in the testing set that have their distances to the training set smaller than the first quartile (Q1), from Q1 to the second quartile (Q2), from Q2 to the third quartile (Q3), and larger than Q3 of the sorted list. Figure 3 shows the average Pearson correlation scores with these distances of three models including DeepCE, vanilla neural network, and kNN. From this figure, we can see the same pattern for all models that the prediction performances are higher when the experiments in the testing set are more similar to their nearest neighbor experiments on the training set. We also recognize that DeepCE achieves better performances than vanilla neural network and kNN for all distance categories, especially for experiments that have their distances to the training set smaller than Q1.

**Figure 3.**
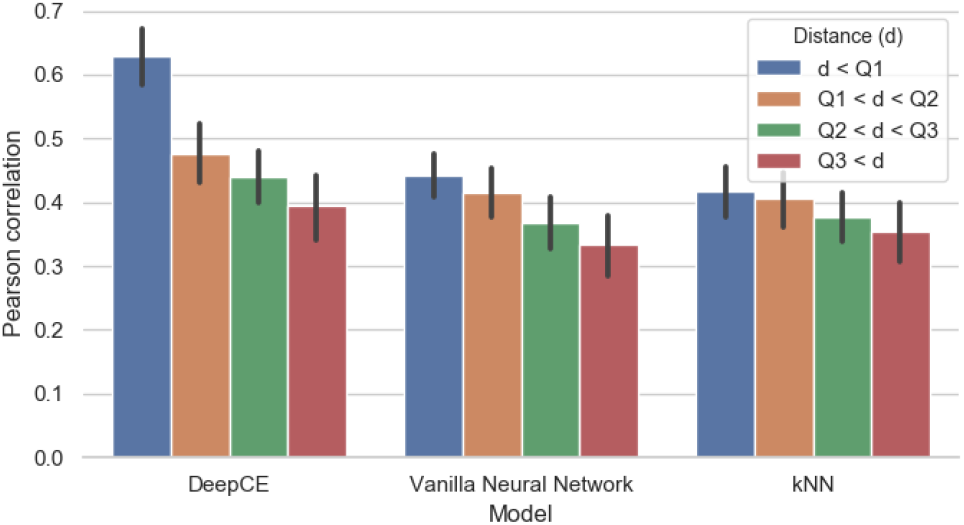
Performances of DeepCE, vanilla neural network, and kNN with different distances among chemicals in the training and testing sets

### Data quality has a significant impact on prediction performance

Besides sparseness problem, L1000 dataset also includes many unreliable gene expression profiles. To investigate the impact of noisy profiles on the prediction performances of our models, we train two baseline models including neural network and kNN on different training sets generated by filtering unreliable gene expression profiles with different average Pearson correlation (APC) thresholds varying from −1 (original training set) to 0.7 (high-quality training set). Chemical feature used in this experiment is PubChem fingerprint.

As shown in Figure 4, all models have the same pattern. Starting at the threshold of 0.1, they achieve better performances on the testing set when the threshold is higher and the best setting is training our models on the high-quality training set (i.e. Pearson correlation of 0.3923 for vanilla neural network and 0.3903 for kNN). For training on the original training set and other training sets generated by filtering unreliable experiments with thresholds <0.1, the ground-truth and predicted gene expression profiles are uncorrelated showing the randomness of the model predictions. These results indicate that unreliable data has a severely negative impact on prediction performances and removing this part from the dataset is necessary for achieving good performances.

**Figure 4.**
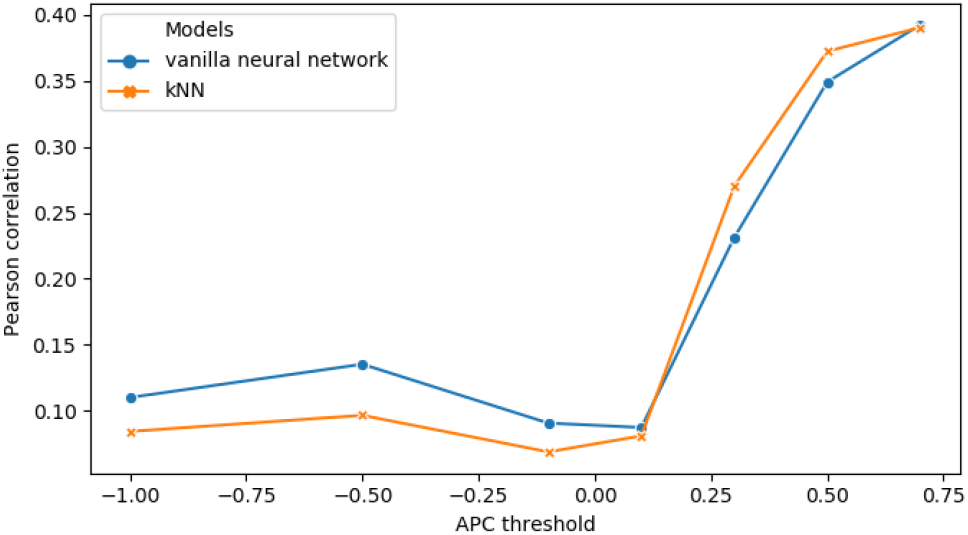
Pearson correlation scores of vanilla neural network and kNN trained on training sets generated by filtering unreliable experiments with different APC thresholds

### A novel data augmentation method improves the model performance

We propose the data augmentation method (described in detail in Supplementary Algorithm 2) to effectively exploit useful information from unreliable gene expression profiles. In this experiment, we evaluate the impact of this method on our models. In particular, DeepCE trained on high-quality training set are used to generate gene expression profiles and the threshold for selecting bio-replicate profiles is 0.5 which is similar to the performance of DeepCE. The statistics of this augmented training set are shown in Supplementary Table 1.

The experimental results for training vanilla neural network, kNN, and DeepCE on the augmented training set are shown in Table 1 and Table 2. We can see that the performances of all models trained on this augmented training set are improved in most cases. For example, the Pearson correlation of DeepCE is increase from 0.4907 to 0.5014 (paired t-test, *p – value* < 0.05). These results indicate that information extracted from unreliable gene expression profiles by our data augmentation method is effective for gene expression prediction.

### Selection of chemical feature affects model performance

In this experiment, we investigate the effectiveness of several chemical feature representations for our models. Models used in this experiment are vanilla neural network for PubChem, ECFP fingerprints, our proposed drug-target features, and LTIP, and DeepCE model without interaction component for neural fingerprint. These models are trained on the high-quality training set. We also create random chemical features by generating random binary vectors whose size is similar to PubChem fingerprint from discrete uniform distribution.

Table 1 (vanilla neural network) and Table 2 (DeepCE) show the performances measured by Pearson correlation of these models with different chemical feature representations. First, chemical features achieve much better performances than the random feature, indicating that chemical features capture important information about chemicals which is useful for predicting gene expression profiles. Second, DeepCE which uses neural fingerprint achieves the Pearson correlation of 0.4418 which is the best performance compared to other settings (paired t-test, *P – value* < 4.89 × 10^−5^). For other chemical features, biological-based features including drugtarget feature and LTIP achieves better performances than chemical-based features including PubChem and ECFP fingerprints. All of these observations are verified by the paired t-tests with *P – values* < 0.01. In fact, most of the *P – values* are much less than 0.01.

### DeepCE is effective in predictive down-stream tasks

In this section, we design an experiment to answer a question about whether these predicted gene expression profiles can provide added values for downstream prediction tasks, especially in the case that original gene expression profiles in L1000 dataset are unreliable. We first extract gene expression profiles of chemicals that do not have reliable experiments in L1000 dataset (original feature set) as well as use DeepCE model trained on high quality training set to generate gene expression profiles for these drugs (predicted feature set). We then use these sets as the features for drugs to train classification models for two tasks: ATC code and drug-target predictions. The details of constructing these datasets are presented in Supplementary Notes and Supplementary Table 4. Finally, we train 4 popular classification models including logistic regression (LR), support vector machine (SVM), k-nearest neighbor (kNN), and decision tree (DT) using 14 different versions of chemical features (7 cell-specific features for each original and predicted feature sets) for 14 binary classification tasks (i.e. 10 ATC codes and 4 drug-targets). For each experiment setting, we use 5-fold cross-validation and report the average results.

The differences in AUC between training classification models with predicted and original feature sets for drug-target and ATC prediction tasks are shown in Figure 5. The improvements in AUC when using predicted features instead of original features are recognized in all cell-specific profiles (Figure 5a), all classification models (Figure 5b), 8/10 ATC codes (Figure 5c), and 3/4 drug-targets (Figure 5d), and these improvements are significant (paired t-test, *P – value* < 4.87 × 10^−5^). The details of AUC scores for predicted and original features for each setting (i.e. per model, cell line, ATC code, and drug-target) are shown in Supplementary Table 5. These results indicate that we can substitute unreliable gene expression profiles in L1000 dataset with gene expression profiles generated from DeepCE to achieve better performances on downstream prediction tasks.

**Figure 5.**
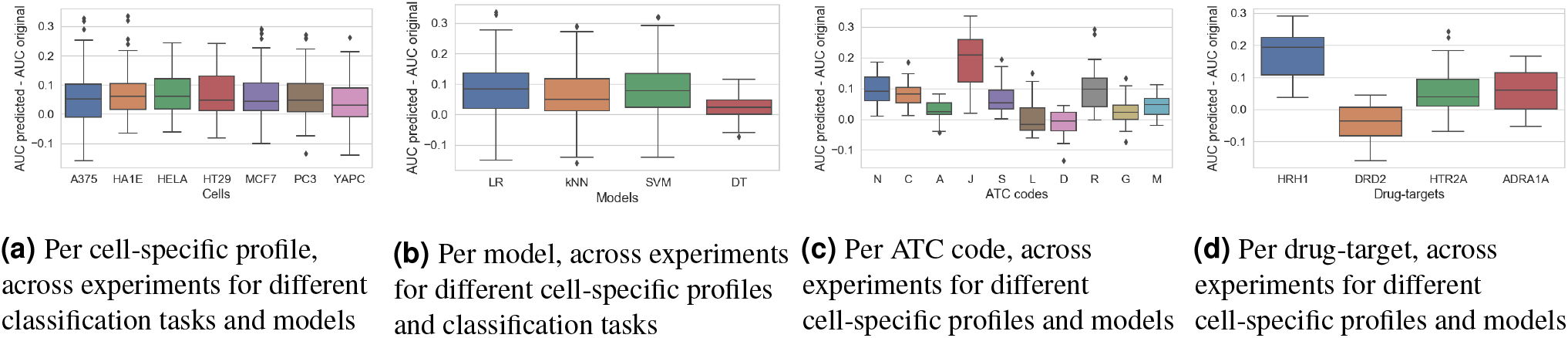
Improvement of predicted profiles over original profiles in AUC

### Drug repurposing for COVID-19

To further demonstrate the value of DeepCE, we use chemical-induced gene expression profile to discover potential drugs for COVID-19 treatment. In particular, we first use trained DeepCE on the high-quality part of L1000 dataset to generate predicted gene expression profiles for all of 11179 drugs in Drugbank database at the largest chemical dose size. We then screen drugs in Drugbank by computing Spearman’s rank-order correlation scores between their gene expression profiles with the patient gene expression profile (see Method section) and select drugs that give the most negative scores as the potential drugs. Here, we incorporate the gene expression profiles of A549 - the cancerous lung tissue - beside the main 7 cell lines in the high-quality dataset. Besides the predicted profiles, we also include the gene expression profiles extracted from the high-quality part of L1000 dataset. For each cell line, we extract the top 100 drugs that have the most negative correlation scores with the patient profile as the potential drugs. Finally, we output top 15 drugs that are potential drugs for COVID-19 treatment at all cell lines as the result of our screening process.

As shown in Table 3, among the 15 drugs we identified, 9 drugs are antiviral drugs and 7 of them are used in treating Hepatitis C as NS5A inhibitor. Especially, two of Hepatitis C treatment, Elbasvir and Velpatasvir, have been shown as potential candidates for COVID-19 treatment by using other approaches^37–39^. Moreover, two drugs shows anti-inflammatory or immune regulating function, and have the potential in regulate immune response under COVID-19 infection. Laniquidar can suppress the function of P-glycoprotein 1 and affect transportation of immunosuppressive agents. AMG-487 targets Chemokine receptor CXCR3, which can regulate leukocyte trafficking. It is noted that all potential drugs here are not available in L1000 dataset, showing the effectiveness of DeepCE for phenotype compound screening.

**Table 3.**
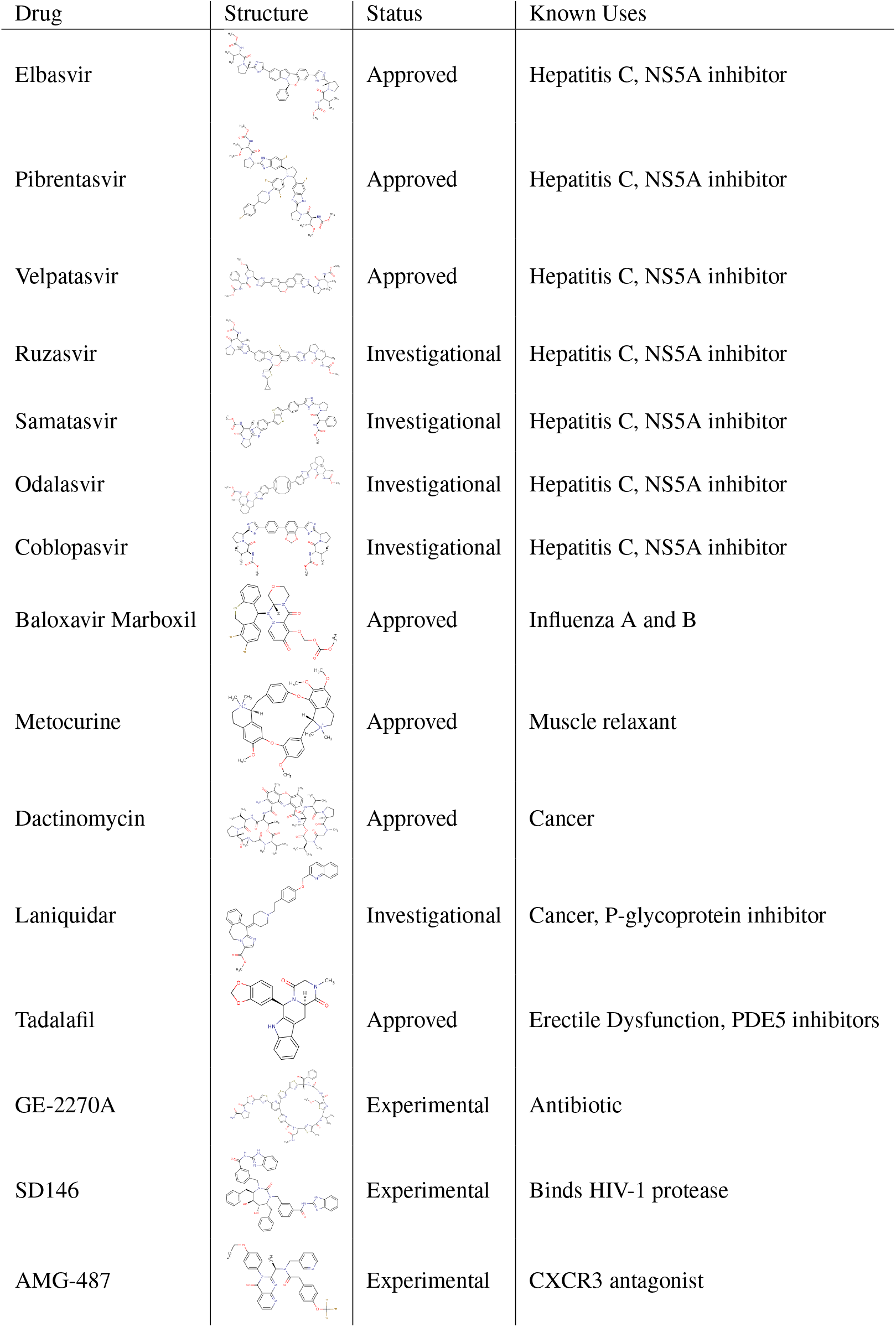
The chemical structures, status, and known uses of potential drugs for COVID-19 treatment (i.e. drugs appeared in top 100 drugs for all 8 cell lines when comparing their cell-specific predicted gene expression profiles with the patient profile by Spearman’s correlation.

| Drug | Structure | Status | Known Uses |
| --- | --- | --- | --- |
| Elbasvir |  | Approved | Hepatitis C, NS5A inhibitor |
| Pibrentasvir |  | Approved | Hepatitis C, NS5A inhibitor |
| Velpatasvir |  | Approved | Hepatitis C, NS5A inhibitor |
| Ruzasvir |  | Investigational | Hepatitis C, NS5A inhibitor |
| Samatasvir |  | Investigational | Hepatitis C, NS5A inhibitor |
| Odalasvir |  | Investigational | Hepatitis C, NS5A inhibitor |
| Coblopasvir |  | Investigational | Hepatitis C, NS5A inhibitor |
| Baloxavir Marboxil |  | Approved | Influenza A and B |
| Metocurine |  | Approved | Muscle relaxant |
| Dactinomycin |  | Approved | Cancer |
| Laniquidar |  | Investigational | Cancer, P-glycoprotein inhibitor |
| Tadalafil |  | Approved | Erectile Dysfunction, PDE5 inhibitors |
| GE-2270A |  | Experimental | Antibiotic |
| SD146 |  | Experimental | Binds HIV-1 protease |
| AMG-487 |  | Experimental | CXCR3 antagonist |

## Conclusion

Deep learning has attracted a great attention in drug discovery. Past and existing efforts mainly focus on accelerating compound screening against a single target^40^. However, such one-drug-one-gene paradigm is proved to be less successful in tracking complex diseases. A systematic compound screening approach, which both takes information on biological system into consideration and uses chemical-induced systematic response as readouts, will provide new opportunities on discovering safe and effective therapeutics that module the biological system. In this study, we have proposed DeepCE - a novel and robust neural network-based model for predicting chemical-induced gene expression profiles from chemical and biological objects, especially in *de novo* chemical setting. Our model achieves state-of-the-art results of predicting gene expression profiles compared to other models not only in *de novo* chemical setting but also in the traditional setting. In addition, we have addressed the unreliable measurement problem of L1000 dataset by introducing the data augmentation method to effectively exploit useful information from unreliable gene expression profiles to improve the prediction performances of our models. Furthermore, the downstream prediction task evaluation shows that training classification models with gene expression profiles generated from DeepCE achieves better performances than training them with unreliable gene expression profiles in L1000 dataset, indicating the added values of DeepCE for downstream prediction. Finally, DeepCE is shown to be effective in the challenge and urgent problem, finding treatment for COVID-19, by *in silico* screening all chemical compounds in Drugbank against COVID-19 patient clinical phenotypes (i.e. comparing chemical-induced gene expression profiles generated from DeepCE with the patient profiles). In summary, DeepCE could be a powerful tool for phenotype-based compound screening.

## Data availability

Chemical-induced gene expression dataset used in our study, gene expression profiles generated from DeepCE for all drugs in Drugbank, source code and usage instructions are available at https://github.com/pth1993/DeepCE.

## Acknowledgements

This work was supported by the National Institute of General Medical Sciences (NIGMS) [R01GM122845 (LX)]; National Institute on Aging of the National Institute of Health (NIH) [R01AD057555 (LX)].

## Author contributions

LX and PZ conceived the project. THP, YQ and JZ extracted and preprocessed the data. THP and PZ developed the method. THP conducted the experiments. THP, YQ, and JZ analyzed the results. THP, YQ, LX, and PZ wrote the manuscript. All authors read and approved the final manuscript.

## References

1. Dudley, J. T., Deshpande, T. & Butte, A. J. Exploiting drug–disease relationships for computational drug repositioning. Briefings bioinformatics 12, 303–311 (2011).

2. Lamb, J. et al. The connectivity map: using geneexpression signatures to connect small molecules, genes, and disease. science 313, 1929–1935 (2006).

3. Hu, G. & Agarwal, P. Human disease-drug network based on genomic expression profiles. PloS one 4 (2009).

4. Kosaka, T. et al. Identification of drug candidate against prostate cancer from the aspect of somatic cell reprogramming. Cancer science 104, 1017–1026 (2013).

5. Wei, G. et al. Gene expression-based chemical genomics identifies rapamycin as a modulator of mcl1 and glucocorticoid resistance. Cancer cell 10, 331–342 (2006).

6. Hassane, D. C. et al. Discovery of agents that eradicate leukemia stem cells using an in silico screen of public gene expression data. Blood, The J. Am. Soc. Hematol. 111, 5654–5662 (2008).

7. Stegmaier, K. et al. Gene expression–based high-throughput screening (ge-hts) and application to leukemia differentiation. Nat. genetics 36, 257–263 (2004).

8. Lamb, J. The connectivity map: a new tool for biomedical research. Nat. reviews cancer 7, 54–60 (2007).

9. Subramanian, A. et al. A next generation connectivity map: L1000 platform and the first 1,000,000 profiles. Cell 171, 1437–1452 (2017).

10. Duan, Q. et al. L1000cds 2: Lincs l1000 characteristic direction signatures search engine. NPJ systems biology applications 2, 1–12 (2016).

11. Iwata, M., Sawada, R., Iwata, H., Kotera, M. & Yamanishi, Y. Elucidating the modes of action for bioactive compounds in a cell-specific manner by large-scale chemically-induced transcriptomics. Sci. reports 7, 40164 (2017).

12. Méndez-Lucio, O., Baillif, B., Clevert, D.-A., Rouquié, D. & Wichard, J. De novo generation of hit-like molecules from gene expression signatures using artificial intelligence. Nat. Commun. 11, 1–10 (2020).

13. Sterling, T. & Irwin, J. J. Zinc 15–ligand discovery for everyone. J. chemical information modeling 55, 2324–2337 (2015).

14. Troyanskaya, O. et al. Missing value estimation methods for dna microarrays. Bioinformatics 17, 520–525 (2001).

15. Bø, T. H., Dysvik, B. & Jonassen, I. Lsimpute: accurate estimation of missing values in microarray data with least squares methods. Nucleic acids research 32, e34–e34 (2004).

16. Kim, H., Golub, G. H. & Park, H. Missing value estimation for dna microarray gene expression data: local least squares imputation. Bioinformatics 21, 187–198 (2005).

17. Cai, Z., Heydari, M. & Lin, G. Iterated local least squares microarray missing value imputation. J. bioinformatics computational biology 4, 935–957 (2006).

18. Oba, S. et al. A bayesian missing value estimation method for gene expression profile data. Bioinformatics 19, 2088–2096 (2003).

19. Ouyang, M., Welsh, W. J. & Georgopoulos, P. Gaussian mixture clustering and imputation of microarray data. Bioinformatics 20, 917–923 (2004).

20. Wang, X., Li, A., Jiang, Z. & Feng, H. Missing value estimation for dna microarray gene expression data by support vector regression imputation and orthogonal coding scheme. BMC bioinformatics 7, 32 (2006).

21. Lagunin, A., Ivanov, S., Rudik, A., Filimonov, D. & Poroikov, V. Digep-pred: web service for in silico prediction of drug-induced gene expression profiles based on structural formula. Bioinformatics 29, 2062–2063 (2013).

22. Perros, I. et al. Polyadic regression and its application to chemogenomics. In Proceedings of the 2017 SIAM International Conference on Data Mining, 72–80 (SIAM, 2017).

23. Hodos, R. et al. Cell-specific prediction and application of drug-induced gene expression profiles. In Pac. Symp. Biocomput, vol. 23, 32–43 (World Scientific, 2018).

24. Iwata, M. et al. Predicting drug-induced transcriptome responses of a wide range of human cell lines by a novel tensor-train decomposition algorithm. Bioinformatics 35, i191–i199 (2019).

25. Liu, C. et al. Compound signature detection on lincs l1000 big data. Mol. BioSystems 11, 714–722 (2015).

26. Li, Z., Li, J. & Yu, P. l1kdeconv: an r package for peak calling analysis with lincs l1000 data. BMC bioinformatics 18, 356 (2017).

27. Qiu, Y., Lu, T., Lim, H. & Xie, L. A Bayesian approach to accurate and robust signature detection on LINCS L1000 data. Bioinformatics (2020).

28. Szklarczyk, D. et al. String v11: protein–protein association networks with increased coverage, supporting functional discovery in genome-wide experimental datasets. Nucleic acids research 47, D607–D613 (2019).

29. Wishart, D. S. et al. Drugbank: a comprehensive resource for in silico drug discovery and exploration. Nucleic acids research 34, D668–D672 (2006).

30. Blanco-Melo, D. et al. Sars-cov-2 launches a unique transcriptional signature from in vitro, ex vivo, and in vivo systems. bioRxiv (2020).

31. Love, M. I., Huber, W. & Anders, S. Moderated estimation of fold change and dispersion for rna-seq data with deseq2. Genome biology 15, 550 (2014).

32. Duvenaud, D. K. et al. Convolutional networks on graphs for learning molecular fingerprints. In Advances in neural information processing systems, 2224–2232 (2015).

33. Vaswani, A. et al. Attention is all you need. In Advances in neural information processing systems, 5998–6008 (2017).

34. Kotlyar, M., Fuhrman, S., Ableson, A. & Somogyi, R. Spearman correlation identifies statistically significant gene expression clusters in spinal cord development and injury. Neurochem. research 27, 1133–1140 (2002).

35. Allison, D. B., Page, G. P., Beasley, T. M. & Edwards, J. W. DNA microarrays and related genomics techniques: design, analysis, and interpretation of experiments (CRC Press, 2005).

36. Ayed, M., Lim, H. & Xie, L. Biological representation of chemicals using latent target interaction profile. BMC bioinformatics 20, 674 (2019).

37. Mevada, V. et al. Drug repurposing of approved drugs elbasvir, ledipasvir, paritaprevir, velpatasvir, antrafenine and ergotamine for combating covid19. (2020).

38. Wang, J. Fast identification of possible drug treatment of coronavirus disease-19 (covid-19) through computational drug repurposing study. J. Chem. Inf. Model. (2020).

39. Shah, B., Modi, P. & Sagar, S. R. In silico studies on therapeutic agents for covid-19: Drug repurposing approach. Life Sci. 252, 117652 (2020).

40. Zhavoronkov, A. et al. Deep learning enables rapid identification of potent ddr1 kinase inhibitors. Nat. biotechnology 37, 1038–1040 (2019).

