## supplementary information for "A deep learning framework for high-throughput mechanism-driven phenotype compound screening"

### ABSTRACT

This supplementary document includes supplementary notes, algorithms, figures, and tables that support the manuscript "A deep learning framework for high-throughput mechanism-driven phenotype compound screening".

### Supplementary Notes

#### Data processing

**L1000 data processing for *de novo* chemical and imputation settings** In this section, we present the method of constructing training, development, and testing sets from Bayesian-based peak deconvolution L1000 dataset which has been shown to generate more robust z-score profiles from L1000 assay data, and therefore, gives better representation for perturbagens. To achieve that, they propose a peak deconvolution algorithm based on Bayes' theorem that gives unbiased likelihood estimations for peak locations and they characterize the peaks with probability-based z-scores.

Bayesian-based peak deconvolution L1000 dataset consists of three levels of the data (corresponding to level 2, 4, and 5 of the original L1000 dataset): level 2 data presents the marginal distributions of peak locations computed from Bayesian-based method, level 4 data presents the z-score of each gene for each bio-replicates by comparing the probability distribution for each gene with its background distribution, and level 5 data combines z-score profiles from bio-replicates into one signature by weighted average. In our study, we conduct experiments on level 5 data and use level 4 data for filtering out unreliable experiments in level 5 data. In particular, average Pearson correlation (APC) score for each experiment in level 5 data is calculated from Pearson correlation scores among its bio-replicates' gene expression profiles in level 4 data and experiments that have APC scores less than a threshold are considered as unreliable experiments and filtered out. In our setting, this threshold is set at 0.7. The APC score density function and cumulative distribution of L1000 data are shown in Supplementary Figure 2. Level 4 data is also used in our data augmentation method described in the following sections. From this filtered dataset, we select the gene signatures measured after 24h of the 7 most popular cell lines (MCF7, A375, HT29, PC3, HA1E, YAPC, HELA) and 6 most popular dose sizes ($0.04\,\mu m$, $0.12\,\mu m$, $0.37\,\mu m$, $1.11\,\mu m$, $3.33\,\mu m$, $10.0\,\mu m$) to create the dataset used in our experiments. We split this dataset into training, development, and testing sets w.r.t. chemical by the ratio $0.6 : 0.2 : 0.2$ respectively. This training set is called high-quality training set. We also create the training set without removing unreliable experiments and call it as original training set. The statistics of these training, development, and testing sets are shown in Supplementary Table 1.

Besides constructing training, development, and testing sets for *de novo* chemical setting, we also generate these set for traditional imputation setting from high-quality dataset. In particular, new training, development, and testing sets are constructed from high-quality dataset with the same ratio as *de novo* chemical setting, but at this time, we split this dataset by experiment instead of chemical. Therefore, chemicals in the testing set can appear in the training set. The statistics of the training, development, and testing sets for traditional imputation setting are shown in Supplementary Table 2

**Data processing for ATC code and drug-target prediction** We extract gene expression profiles of chemicals that do not have reliable experiments in L1000 dataset (i.e. 1258 chemicals) and call them a original feature set. We only extract gene expression profiles induced by chemicals with dose size of 10 um because gene expression profiles induced by the larger amount of chemicals can differentiate chemicals better than them induced by the smaller amount of chemicals. Note that, each chemical compound is experimented on 7 cell lines so it is represented by 7 gene expression profiles. Similarly, we use DeepCE trained on the high-quality training set to generate gene expression profiles for these chemicals and call them a predicted feature set. Next, we extract the ATC codes and drug-targets of these chemicals from Drugbank database to create labels for these chemicals. We also filter out labels that have low frequencies in the dataset (i.e. $< 3\%$ of the number of chemicals) to avoid unreliable evaluation. After that, 10 ATC codes ((i.e. *N, C, A, J, S, L, D, R, G, M*) and 4 drug-targets (HRH1, DRD2, HTR2A, ADRA1A) are selected to construct 14 binary classification datasets. The details of these datasets are shown in Supplementary Table 4.

#### Feature engineering

Our models predict gene expression values based on tuples of chemical and biological objects including chemical compounds, cell lines, dose sizes, and L1000 genes. In the following paragraphs, we present the way to transform these chemical and biological objects into numerical representations that can be put into our models.

**Chemical fingerprints** The canonical SMILES strings which are the raw representation of chemicals are transformed to the chemical fingerprints using the open-source cheminformatics software RCDK[1] (i.e. *get.fingerprint()* function). chemical fingerprints are binary (bit) vectors that represent the presence or absence of particular substructures in chemicals. In our settings, we experiment with PubChem and circular (ECFP6) fingerprints which have lengths (i.e. number of substructures) of 881 and 1024 respectively. We also use neural fingerprints which are continuous vectors to represent chemicals. The neural fingerprints are described in detail later.

**Drug-target features** Besides using molecule structure information, bioactivity information of chemicals (i.e. drug-target interaction knowledge) can be used to represent chemicals. In particular, vector representations for chemicals are calculated from drug-target interaction and human protein-protein interaction networks extracted from Drugbank and STRING databases respectively. To increase the quality of generated vectors, all edges (interactions) in the human protein-protein interaction network have their combined scores of less than 700 are removed. If chemical compound interacts with L1000 genes (i.e. L1000 genes are 1-hop neighborhoods of chemical compound in the interaction network), values at indexes corresponding to these genes are set at 1.0. If L1000 genes interact with targets of chemicals (i.e. L1000 genes are 2-hop neighborhoods of chemical compound in interaction network), their values are set at 0.1. Similarly, the values of L1000 genes which are 3-hop neighborhoods of chemical compound are set at 0.01. The values of remaining L1000 genes are set at 0. The generated vectors are normalized to have Euclidean norm to be 1. The length of these vectors is 978 which is equal to the number of L1000 genes. Besides our proposed drug-target interaction-based representation, we also experiment with another chemical representation using drug-target interaction information named latent target interaction profile (LTIP) which has been shown to be an effective representation for chemical compound in many bioactivity prediction tasks, especially for the setting that the biological information is important for prediction[2]. In particular, LTIP maps chemicals into low dimensional continuous vectors which are embeddings of nodes in the drug-target interaction network (i.e. bipartite graph where drugs and targets are nodes, and their interactions are edges).

**Gene features** Vector representations for L1000 genes are generated from the human protein-protein interaction network extracted from STRING database using node2vec method[3]. The length of these vectors is set at 128.

**Cell line and chemical dose size features** We represent cell lines and chemical dose sizes using one-hot encoding. The lengths of these vectors are 7 and 6 corresponding to the numbers of cell lines and chemical dose sizes used in our experiments respectively.

#### Implementation details

All neural network-based models including DeepCE, vanilla neural network, and linear models are implemented with Pytorch[4]. The training process lasts at most 100 epochs for all training sets and all models. We use Adam optimizer[5] with the learning rate of 0.0005 and the batch size is set at 16. The development set is used for tuning hyperparameters and stopping the training process. kNN model is implemented by Python 3 while we use authors' implementation for TT-WOPT. Binary classification models used in downstream task evaluation including logistic regression, support vector machine and kNN are implemented by scikit-learn package[6]. All experiments are conducted on Intel Xeon E5-2680 v4 processor with 128 GB of RAM and Tesla P100 with 16 GB of VRAM and are repeated three times with different random seeds.

### Supplementary Tables

| | #chemicals | Cell lines (#gene expression profile) | | | | | | | Chemical doses (#gene expression profile) | | | | | |
|---|---|---|---|---|---|---|---|---|---|---|---|---|---|---|
| | | A375 | HA1E | HELA | HT29 | MCF7 | PC3 | YAPC | 0.04 um | 0.12 um | 0.37 um | 1.11 um | 3.33 um | 10.00 um |
| Train (Original) | 1553 | 9225 | 9233 | 8443 | 9243 | 9240 | 9247 | 8440 | 10587 | 10504 | 10526 | 10512 | 10502 | 10440 |
| Train (High-quality) | 284 | 397 | 296 | 209 | 287 | 250 | 279 | 177 | 136 | 175 | 227 | 322 | 402 | 633 |
| Train (Augmented) | 626 | 707 | 953 | 399 | 552 | 519 | 514 | 321 | 327 | 414 | 497 | 639 | 817 | 1271 |
| Dev | 92 | 120 | 81 | 50 | 89 | 77 | 69 | 58 | 40 | 57 | 72 | 93 | 89 | 193 |
| Test | 92 | 121 | 69 | 39 | 82 | 61 | 72 | 52 | 33 | 48 | 57 | 83 | 91 | 184 |

**Supplementary Table 1.** Statistics of (original/high-quality/augments) training, development, and testing sets generated from Bayesian-based peak deconvolution L1000 dataset for *de novo* chemical setting

| | #chemicals | Cell lines (#gene expression profile) | | | | | | | Chemical doses (#gene expression profile) | | | | | |
|---|---|---|---|---|---|---|---|---|---|---|---|---|---|---|
| | | A375 | HA1E | HELA | HT29 | MCF7 | PC3 | YAPC | 0.04 um | 0.12 um | 0.37 um | 1.11 um | 3.33 um | 10.00 um |
| Train | 394 | 396 | 258 | 177 | 288 | 229 | 251 | 162 | 133 | 169 | 235 | 292 | 332 | 600 |
| Dev | 268 | 132 | 89 | 54 | 90 | 76 | 82 | 64 | 44 | 51 | 61 | 99 | 129 | 203 |
| Test | 257 | 110 | 99 | 67 | 80 | 83 | 87 | 61 | 32 | 60 | 60 | 107 | 121 | 207 |

**Supplementary Table 2.** Statistics of training, development, and testing sets generated from Bayesian-based peak deconvolution L1000 dataset for traditional imputation setting

| | Hyperparameter | Value |
|---|---|---|
| Training | Batch size | 16 |
| | Learning rate | 0.0002 |
| | Optimizer | Adam |
| | Maximum number of epochs | 100 |
| | Loss function | MSE |
| | Initializer | Xavier Uniform |
| Feature mapping | Atom feature size | 62 |
| | Bond feature size | 6 |
| | Convolution size | 16 |
| | Number of convolutional layers | 2 |
| | Neural fingerprint size | 128 |
| | Chemical dose feature size | 6 |
| | Cell line feature size | 7 |
| | L1000 gene feature size | 128 |
| | Chemical dose hidden size | 4 |
| | Cell line hidden size | 4 |
| | L1000 gene hidden size | 128 |
| Interaction component | Number of attention heads | 4 |
| | Number of attention layers | 2 |
| | Attention hidden size | 512 |
| | Batch normalization | yes |
| | Dropout | 0.1 |
| Prediction component | Number of feed-forward layers | 2 |
| | Feed-forward layer sizes | 128, 978 |
| | Activation function | ReLU |

**Supplementary Table 3.** Hyperparameter used in DeepCE model.

| ATC code | Frequency | ATC code | Frequency |
|---|---|---|---|
| N | 143 | L | 57 |
| C | 124 | D | 54 |
| A | 84 | R | 53 |
| J | 80 | G | 48 |
| S | 62 | M | 43 |

**(a)** Frequencies of ATC codes across all low-quality drugs

| Drug-target | Frequency |
|---|---|
| HRH1 | 47 |
| DRD2 | 45 |
| HTR2A | 42 |
| ADRA1A | 40 |

**(b)** Frequencies of drug-targets across all low-quality drugs

**Supplementary Table 4.** Data statistics for downstream task evaluation

| | Original feature | Predicted feature |
|---|---|---|
| A375 | 0.5712 | 0.6292 |
| HA1E | 0.5441 | 0.6173 |
| HELA | 0.5527 | 0.6262 |
| HT29 | 0.5522 | 0.6182 |
| MCF7 | 0.5518 | 0.6203 |
| PC3 | 0.5586 | 0.6206 |
| YAPC | 0.5805 | 0.6191 |

**(a)** Per cell-specific profile, across experiments for different classification tasks and models

| | Original feature | Predicted feature |
|---|---|---|
| N | 0.5534 | 0.6511 |
| C | 0.5260 | 0.6077 |
| A | 0.5047 | 0.5312 |
| J | 0.5199 | 0.7116 |
| S | 0.5678 | 0.6385 |
| L | 0.5153 | 0.5219 |
| D | 0.5923 | 0.5806 |
| R | 0.4969 | 0.5955 |
| G | 0.5436 | 0.5698 |
| M | 0.5436 | 0.5890 |

**(b)** Per ATC code, across experiments for different cell-specific profiles and models

| | Original feature | Predicted feature |
|---|---|---|
| HRH1 | 0.6015 | 0.7746 |
| DRD2 | 0.6219 | 0.5792 |
| HTR2A | 0.6538 | 0.7115 |
| ADRA1A | 0.5813 | 0.6399 |

**(c)** Per drug-target, across experiments for different cell-specific profiles and models

| | Original feature | Predicted feature |
|---|---|---|
| LR | 0.5958 | 0.6809 |
| kNN | 0.5471 | 0.6110 |
| SVM | 0.5850 | 0.6655 |
| DT | 0.5070 | 0.5288 |

**(d)** Per model, across experiments for different cell-specific profiles and classification tasks

**Supplementary Table 5.** AUC of experiments with original and predicted gene expression profiles

### Supplementary Algorithms

**Algorithm 1:** Pseudo-code of GCN

**Input:** Chemical graph = $(V,E)$, radius $R$, hidden weights $(H_1^1,...,H_R^5),(U_1,...,U_l),(W_1,...,W_l)$
**Output:** $v_R^{(1)},...,v_R^{(|V|)}$
**for** *l = 1 to R* **do**
  **for** *i = 1 to —V—* **do**
    $V_{neighbor}, E_{neighbor} \leftarrow Neighbors(\mathbf{v}^{(i)})$;
    $\mathbf{v}_l^{(i)} \leftarrow \sum_{\mathbf{v}^{(j)} \in V_{neighbor}} \mathbf{v}_{l-1}^{(j)}$;
    $\mathbf{e}_l^{(i)} \leftarrow \sum_{\mathbf{e}^{(j)} \in E_{neighbor}} \mathbf{e}_0^{(j)}$;
    $\mathbf{v}_l^{(i)} \leftarrow concat(\mathbf{v}_l^{(i)}, \mathbf{e}_l^{(i)})$;
    $\mathbf{v}_l^{(i)} \leftarrow ReLU(\mathbf{v}_{l-1}^{(i)}\mathbf{U}_l + (\mathbf{v}_l^{(i)}\mathbf{W}_l)$;
    $\mathbf{v}_l^{(i)} \leftarrow softmax(\mathbf{v}_l^{(i)}\mathbf{H}_l^{|V_{neighbor}|})$
  **end**
**end**

**Algorithm 2:** Pseudo-code of data augmentation method

**Input:** High-quality training set $D_{high}^{LV5}$, level 4 low-quality training set $D_{low}^{LV4}$, model $F(\Theta)$, threshold $t$
**Output:** Augmented training set $D_{augment}$
$D_{augment} \leftarrow D_{high}^{LV5}$;
$F(\Theta_{train}) \leftarrow$ train $F(\Theta)$ on $D_{high}^{LV5}$;
$D_{predict} \leftarrow$ predicted gene expression profiles for unreliable experiments by $F(\Theta_{train})$;
**for** *each profile d in* $D_{predict}$ **do**
  $D_{bio} \leftarrow$ set of bio-replicate profiles corresponding to $d$ in $D_{low}^{LV4}$;
  $S \leftarrow$ set of similarity scores of $D_{bio}$ and $d$;
  $d_{max} \leftarrow argmax_{d \in D_{bio}} s_d \in S$;
  **if** $d_{max} \geq t$ **then**
    $D_{augment} \leftarrow D_{augment} \cup d_{max}$
  **end**
**end**

### Supplementary Figures

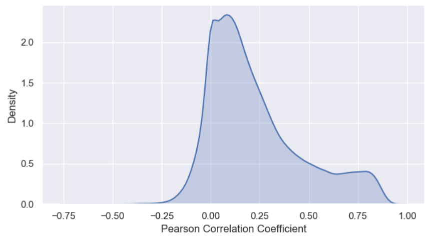


**(a)** Density function of APC score for level 5 data

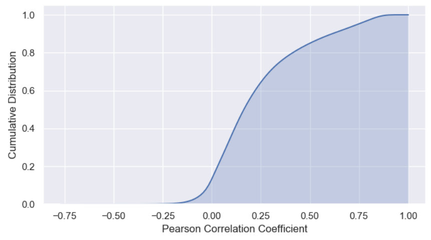


**(b)** Cumulative distribution of APC score for level 5 data

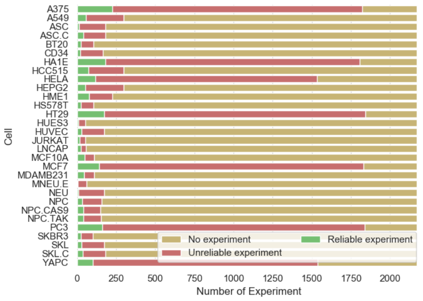


**(c)** Experiment statistics per cell line (time = 24h and dose size = 10 um)

**Supplementary Figure 1.** Statistics of Bayesian-based peak deconvolution L1000 dataset (Experiments that have APC score $< 0.7$ are considered as unreliable experiments).

### Results of DeepCE on the testing test per cell line and chemical dose size

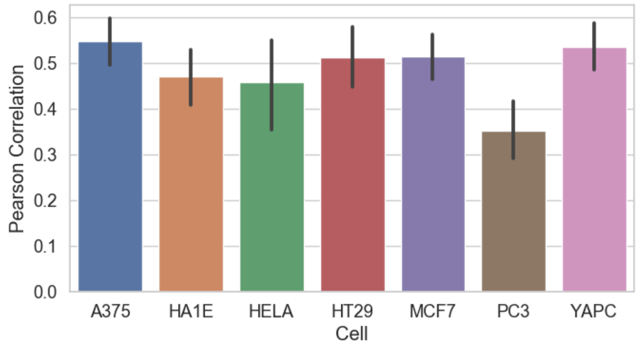


**(a)** Performance of DeepCE model per cell line

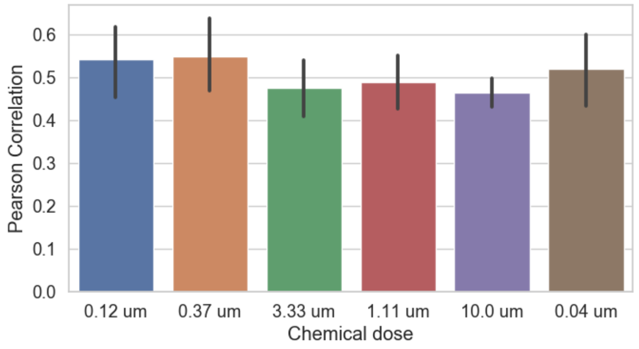


**(b)** Performance of DeepCE model per chemical dose size

**Supplementary Figure 2.** Performance (Pearson correlation) of DeepCE model on the testing set per cell line and chemical dose size

### References


1. Guha, R. Chemical informatics functionality in r. *J. Stat. Softw.* **18**, 1–16 (2007).
2. Ayed, M., Lim, H. & Xie, L. Biological representation of chemicals using latent target interaction profile. *BMC bioinformatics* **20**, 674 (2019).
3. Grover, A. & Leskovec, J. node2vec: Scalable feature learning for networks. In *Proceedings of the 22nd ACM SIGKDD international conference on Knowledge discovery and data mining*, 855–864 (2016).
4. Paszke, A. *et al.* Automatic differentiation in pytorch. In *Proceedings of the 2017 Neural Information Processing Systems Workshop Autodiff* (2017).
5. Kingma, D. P. & Ba, J. Adam: A method for stochastic optimization. In *3rd International Conference on Learning Representations, ICLR 2015, San Diego, CA, USA, May 7-9, 2015, Conference Track Proceedings* (2015).
6. Pedregosa, F. *et al.* Scikit-learn: Machine learning in Python. *J. Mach. Learn. Res.* **12**, 2825–2830 (2011).
